# Kinesin-1 Autoinhibition Tunes Cargo Transport by Motor Ensembles

**DOI:** 10.1101/2025.05.06.652443

**Authors:** Brandon M. Bensel, Samantha B. Previs, Patricia M. Fagnant, Kathleen M. Trybus, Sam Walcott, David M. Warshaw

## Abstract

Intracellular vesicular transport by kinesin-1 motors through numerous 3-dimensional (3D) microtubule (MT) intersections must be regulated to support proper vesicle delivery. Knowing kinesin-1 can be regulated via autoinhibition, does kinesin-1 exhibit autoinhibition on cargo, and could this regulate vesicular transport through 3D MT intersections *in vitro*? To answer this question, we compared liposome transport by ∼10 nearly full-length kinesin-1 motors with KLC bound (KinΔC) versus constitutively active control (K543). In 3D MT intersections, KinΔC-liposomes terminate (48%), go straight (43%), but rarely turn (9%), starkly contrasting K543-liposomes which go straight (57%), turn (31%), but rarely terminate (12%). On single MTs, KinΔC-liposomes have reduced run lengths and detachment forces versus K543-liposomes, suggesting autoinhibition reduces MT engagement, as supported by 3-fold lower KinΔC MT landing rates versus K543, and mechanistic *in silico* modeling. Furthermore, kinesore, a small molecule that overcomes kinesin-1 autoinhibition, restores KinΔC’s MT engagement. Thus, we propose that partial kinesin-1 autoinhibition while cargo-bound may fine-tune cargo delivery to support physiological demands.

**Statement of significance:** Kinesin-1 vesicular transport through the cell’s complex microtubule (MT) network must be spatially and temporally regulated to meet the cell’s physiological demands. With kinesin-1 being autoinhibited, does this form of regulation impact the directional outcomes of vesicles encountering 3-dimensional (3D) MT-MT intersections as they do in cells? To address this in vitro, liposomes transported by kinesin-1 motors (team of ∼10), which exhibit autoinhibition, are challenged with 3D MT intersections and interestingly, preferentially terminate or go straight and rarely turn, while liposomes transported by constitutively active motors preferentially go straight or turn but rarely terminate. Therefore, autoinhibition reduces the number of MT-engaged motors within the team and provides the cell a means of fine-tuning cargo transport to its destination.

## Introduction

Secreted and transmembrane proteins are sorted within the Golgi network into vesicles that are transported by kinesin-1 (KIF5) motors along microtubules (MTs) to their destination on the plasma membrane (1–7). To meet the cell’s physiological demands, vesicular transport and delivery must be regulated spatially and temporally (3, 8). Although multiple mechanisms exist to regulate how kinesin-1 motor teams maneuver vesicular cargo through the complex 3-dimensional (3D) MT network (9–11), the most basic mode of regulation is by activating or inhibiting individual motors (11, 12). Heterotetrameric kinesin-1, consisting of two heavy chains (KHC) and two kinesin light chains (KLC) (13, 14), adopts a “folded”, autoinhibited state with numerous intramolecular interactions along the length of the KHC (1, 15–18). The KLCs further stabilize the inhibited state by forming a 6-helix bundle with the KHC near the hinge 2 region (4, 15–17). Cargo binding to KHC’s C-terminal domains or KLC is thought to disrupt these autoinhibitory interactions, allowing the motor to become “extended” and active (i.e., able to engage the MT) (16). However, once bound to cargo, is kinesin-1 stably active, or is it in an equilibrium between the autoinhibited “folded” and active “extended” states, which could be tuned to optimize cargo transport and delivery by kinesin-1 motor teams (8, 19)?

Here we reconstitute vesicular cargo transport *in vitro* to determine if kinesin-1 autoinhibition affects multimotor transport of physiologically relevant, 350-nm diameter, fluid-like liposomes through suspended 3D MT intersections (20), much like those encountered in cells (21–24). We took advantage of our previously characterized, nearly full-length (1-888 amino acids) kinesin-1 construct (KinΔC) with bound KLC that lacks the so-called IAK region near the C-terminus that is necessary for full inhibition of activity, but retains the hinge region in the coiled coil that allows the molecule to bend and engage in a head–tail interaction (25). Directional outcomes in 3D MT intersections for liposomes transported by teams of ∼10 KinΔC motors were in stark contrast to those of liposomes transported by teams of a constitutively active truncated kinesin-1 lacking KLC (K543) (20). Specifically, KinΔC-liposomes either went straight through or terminated in intersections, but rarely turned onto the intersecting MT. K543-liposomes, preferred to go straight rather than turn, but rarely terminated in the intersection (20). These differences in directional outcomes may be due to the autoinhibition of the cargo-bound KinΔC motors effectively reducing the number of motors available to engage the MT (16, 17, 25). This hypothesis is supported by the observation that cargos transported by KinΔC ensembles along single MTs have shorter run lengths and generate less force against an optical trap compared to K543 ensembles. Treating KinΔC with kinesore, a small molecule that binds KLC and activates kinesin-1 (26), causes the motor to adopt a more “extended” conformation, making KinΔC MT engagement and transport identical to constitutively active K543 motors. To explain why KinΔC autoinhibition modulates liposome transport in 3D MT intersections, we modified our previously published *in silico* mechanistic model of kinesin-1 transport (20) to implicitly include autoinhibition. The model reproduces our observations of liposome transport through 3D MT intersections by KinΔC when modified either by reducing the number of active motors in the ensemble or reducing the maximum rate of MT association, supporting our hypothesis. We propose that kinesin-1 autoinhibition provides the cell with a mechanism to expand and fine tune its dynamic range of potential cargo transport and delivery properties.

## Materials and Methods

### Kinesin motor constructs and purification

Two kinesin motor constructs were used in this study, KinΔC and K543. KinΔC contains the N-terminal 888 amino acids of the mouse KIF5B kinesin heavy chain (accession number: Q61768) followed by a biotin ligase recognition sequence and FLAG tag used for purification, co-expressed with full-length mouse kinesin light chain KLC2 (accession number: BC014845). K543 contains the N-terminal 543 amino acids of the mouse KIF5B kinesin heavy chain (accession number: Q61768) followed by a biotin ligase recognition sequence and FLAG tag used for purification. The biotin ligase recognition sequence used in both constructs is an 88 amino acid sequence from the E. coli biotin carboxyl carrier protein, and is singly biotinylated during expression (27).

Both kinesin constructs were expressed in the baculovirus/Sf9 cell system and purified by affinity chromatography as described previously (20). To express KinΔC, Sf9 cells were infected with two recombinant baculoviruses, one encoding the sequence of the KinΔC heavy chain, the other encoding full-length KLC2. Each baculovirus was used at a ratio of 5 viral particles per cell. Importantly, prior work demonstrated that this strategy yields a 1:1 stoichiometry of KLC to KinΔC heavy chain in the final sample (25, 28). Additional kinesin constructs with an encoded YFP were expressed and purified as described previously for motor counting experiments (20, 29). For motor counting KinΔC, a C-terminal YFP was added to the KLC2. Kinesin aliquots were in 10mM Imidazole (pH = 7.4) with 200 mM NaCl, 55% glycerol, 1 mM DTT, 1μg/ml Leupeptin, and 50 μM MgATP. Aliquots were flash frozen prior to storage at - 80° C.

### Microtubule preparation

Microtubules (MTs) were prepared using a mixture of unlabeled porcine brain tubulin (Cytoskeleton) with rhodamine labelled tubulin (Cytoskeleton) or Alexa-647 labelled tubulin (PurSolutions). For rhodamine labelled microtubules, 20% of the tubulin was labelled, while Alexa-647 microtubules contained 33% labeled tubulin. MTs were assembled and stabilized *in vitro* as described previously (20). The tube containing the assembled MTs was wrapped in foil for storage in a dark drawer, thus minimizing potential photodamage. Polymerized MTs were stable at room temperature for up to one week.

### Single molecule quantum dot motility assays

Single molecule motility assays were performed as described previously (20). Briefly, a kinesin aliquot was rapidly thawed and brought to a final volume of 50 μl with ice cold Buffer 1 (25 mM Imidazole pH = 7.4, 300 mM KCl, 4 mM EGTA, 4 mM MgCl_2_). The protein was then clarified by ultracentrifugation at 392,000 x g for 10 minutes at 4° C, transferred to a fresh tube on ice, then diluted to a final concentration of 100 nM in Buffer 2 (25 mM Imidazole pH = 7.4, 25 mM KCl, 4 mM EGTA, 4 mM MgCl_2_). 2 μl of 100 nM kinesin was mixed with 2 μl of 1 μM Qdot 655 Streptavidin Conjugate (ThermoFisher) and 6 μl Buffer 2 and incubated on ice for 1 hour. The 10:1 ratio of Qdots to kinesin was selected to ensure that experiments are truly in single molecule conditions. These assays were performed identically for K543 and KinΔC (20).

Single molecule experiments were performed on silanized cover glass, which was prepared as described previously (20). Motility chambers were prepared by attaching a 22×22 mm No. 1 cover glass to a silanized 24×60 mm silanized cover glass, separated by 125 μm Mylar shims, using UV-curable adhesive (Norland Optical Adhesive 68). The motility surface was prepared by first coating with 0.8% anti-tubulin antibody (BioRad YL1/2) in BRB-80 and incubating for 5 minutes. The chamber was then washed with Wash Buffer (BRB-80 plus 20 μM paclitaxel) prior to blocking with 5% w/v Pluronic F-127 in BRB-80 for 5 minutes and again washed. Rhodamine-labelled MTs diluted 1:400 in BRB-80 with 20 μM paclitaxel were flowed into the motility chamber and incubated for 10 minutes prior to a final wash with Wash Buffer. Kinesin-Qdot complex was diluted 1:20 in Motility Buffer (Buffer 1 with 2 mM MgATP, 20 μM paclitaxel, 0.5 mg/ml Casein, 0.5% w/v Pluronic F-127, 5 mM creatine phosphate, 0.4 mg/ml creatine phosphokinase, 10 mM DTT, 3.5 mg/ml glucose, 40 μg/ml glucose oxidase, 27 μg/ml catalase) and flowed into the motility chamber. For experiments utilizing the kinesin activator Kinesore (Tocris Bioscence Cat. No. 6664), a Kinesore stock of 50 mM was prepared in DMSO and kept at −20° C. On the day of the experiment, Kinesore was thawed, diluted 1:10 in DMSO, and further diluted 1:100 in Motility Buffer alongside a control buffer containing an equivalent volume of DMSO. Motility videos and still images of MTs were acquired using a custom-built dual-camera TIRF microscope (29, 30) with 532-nm excitation to image MTs and 639-nm excitation to image Qdots. Videos of Qdot motility were acquired at 10 frames per second.

### Data analysis

Qdot motility was analyzed using the Multi Kymograph plugin for ImageJ (31), as published previously (20). Run length and velocity data are plotted as dot plots with the median and upper and lower quartiles overlaid on the plot. To determine MT landing rates, the number of events seen on a single kymograph was divided by the product of the length of the MT in μm and the duration of the movie in minutes. For each condition of interest, multiple kymographs were analyzed from multiple independent videos, and a mean landing rate and standard deviation were determined based on the landing rate observed in each kymograph in each condition.

### Liposome and kinesin-liposome complex preparation

Liposomes used in this study were prepared identically to those described previously (20, 29, 30, 32). The liposomes are stored at room temperature in a dark drawer for up to one week of experiments. This liposome preparation yields 400 μl of liposomes at a final concentration of 10 nM. For fluorescence-based motor counting assays (see below), liposomes were prepared with the fluorescent dye omitted to ensure that no background fluorescence interfered with motor counting.

To prepare kinesin-liposome complexes, either KinΔC or K543 aliquots were thawed and clarified as described above and diluted in Buffer 1 to the desired concentration (250 nM to 1 μM) depending on the kinesin-liposome ratio required for the experiment. 4 μl of diluted kinesin is mixed with 16 μl of liposomes (10 nM) and 20 μl of Buffer 2, and the mixture is incubated on ice for at least one hour. Motor counting experiments were performed as described previously (20, 29, 30). Briefly, KinΔC with YFP-KLCs or a K543 with an N-terminal YFP was clarified, diluted in Buffer 1 as described above (300 nM – 2.5 μM), and incubated with non-fluorescent liposomes. Flow chambers were prepared using plasma-cleaned cover glass. MTs, prepared as described previously (20) with the single change of omitting fluorescently-labelled tubulin, were affixed to the flow cell surface via an anti-tubulin antibody as described above, to promote landing of the kinesin-liposome complex onto the slide surface. YFP-KinΔC-liposome complexes were diluted 1:20 into motility buffer without MgATP, perfused into the chamber, and immediately imaged by TIRF microscopy with 488-nm excitation on a Nikon N-STORM microscope at ∼60 frames per second.

Monitoring the integrated YFP intensity over time for a spot corresponding to a liposome yields the photobleaching transient for that liposome (Fig. S1A). Because the number of fluorescent molecules on each liposome is too large to allow for clear observation of individual photobleaching steps (29), we used a statistical photobleaching technique developed by Nayak and Ruttenberg (33), which we have used and described the analysis routine for in detail previously (20, 29, 30, 32). By performing this analysis across a range of kinesin to liposome ratios, we generated a standard curve of the number of bound kinesins versus the kinesin to liposome ratio in the incubation tube (Fig. S1B and C).

### Multi-motor liposome motility assays and data analysis

Liposome motility assays were performed using DiI labeled liposomes as described previously (20). Briefly, on the day of the experiment, an aliquot of kinesin was thawed, clarified, and diluted as described above. 4 μl of diluted kinesin (concentration range 0.4 to 1.6 μM) was mixed with 16 μl of liposomes (10 nM) and 20 μl of Buffer 2 and set to incubate on ice for at least 1 hour. Motility chambers were prepared as described above for single molecule motility assays. Immediately before imaging, liposomes were diluted 1:20 into Motility Buffer and perfused into the motility chamber prior to imaging by TIRF microscopy on the same custom microscope described above with 532-nm excitation to image MTs and 639-nm excitation to image the DiI liposomes, and motility was recorded at a framerate of 10 frames per second.

### Data analysis

As in the single molecule motility experiments, liposome motility data were analyzed via kymography. Multi-motor runs were much more likely to reach the end of the MT (34–36), so end events were clearly marked in the analysis spreadsheet. Run length and velocity data are shown as dot plots with overlaid median and upper and lower quartiles. End event frequences are shown as bar graphs.

### Optical trapping assays

We use 500-nm lipid-coated silica microspheres as model cargo for both single- and multi-motor trapping experiments, prepared as described previously (20, 30), based on published protocols (37, 38). Briefly, DiI liposomes are prepared but not extruded through the 200 nm pore size extrusion filter (T&T Scientific). Instead, liposomes are washed 3 times by ultracentrifugation (392,000 x g for 10 minutes) to remove excess SH-NaV and resuspended in Buffer 3 (10 mM HEPES pH = 7.2 and 150 mM NaCl). The liposomes are sonicated in an ice bath at low power with 0.5s pulses for a total sonication time of 10 minutes, centrifuged for 10 minutes at 5,000 x g, and transferred to a fresh tube. 50 μl of 500 nm diameter silice microspheres (Duke Standard) are washed with methanol, dried in a speedvac (Rotovap;Eppendorf), and resuspended in 200 μl of Buffer 3. The sonicated liposomes are incubated at 60° C for 2 minutes, mixed with the silica microspheres, and immediately vortexed and briefly sonicated. The mixture is then shaken at room temperature for 1 hour, washed 3 times by low-speed centrifugation and resuspended in PBS (pH = 7.4) prior to storage at 4° C with agitation for up to 3 days.

Optical trapping assays were performed as published previously (20). Briefly, flow chambers were prepared using silanized cover glass. On an experiment day, an aliquot of kinesin was mixed with lipid-coated silica microspheres in 20-fold molar excess, while single molecule force assays were performed with limiting kinesin, as confirmed by screening microspheres to find an incubation ratio where ∼1 of every 10 beads generated force. Flow cells were coated with 0.8% w/v anti-tublin antibody (BioRad YL1/2), blocked with 1 mg/ml Casein (Sigma Aldrich Cat. No. C0406) in Buffer 2, and rhodamine-labeled MTs (1:400 in BRB-80 with 20 μM paclitaxel) were introduced in two sequential flows to align the MTs along a single axis. The motility chamber was then perfused with Motility Buffer prior to adding a small volume of kinesin-bead complex to one end of the chamber. Doing this creates a “bead front” and makes it easier to trap a single lipid-coated silica microsphere at a time. For experiments performed with Kinesore, a Kinesore stock of 50 mM was prepared in DMSO and kept at −20° C. On the day of the experiment, Kinesore was thawed, diluted 1:10 in DMSO, and the resultant Kinesore dilution was further diluted 1:100 in Motility Buffer alongside a control buffer containing an equivalent volume of DMSO. Optical trapping assays were performed on a commercially available optical trap (Lumicks C-Trap System) which is controlled using the supplied software, BlueLake (Lumicks), as published previously (20). The trap stiffness used throughout this study ranged from ∼0.04 to 0.06 pN/nm.

### Data analysis

Optical trapping data were analyzed using a custom-written R script as published previously (20). Detachment force distributions were fit using the fitdistr function of the MASS packing in R (39), using a single Gaussian for single motor experiments and a Gaussian mixture model for multi-motor experiments. Log-likelihood ratio testing was performed to determine if the inclusion of a third Gaussian was statistically justifiable (39).

### 3D MT intersection assay

3D MT intersection assays were performed as described previously (20). Briefly, crossflow motility chambers with two perpendicular flow channels were assembled as published previously (20, 30, 32). Poly-L-Lysine-coated pedestal beads were prepared as described previously (20, 30, 32) and introduced to the flow cell to suspend MTs above the glass surface. Following this, to block the surface in both flow channels, 1 mg/ml BSA in Buffer 2 was infused and incubated for 5 minutes and then washed with Wash Buffer. MTs labelled with Alexa-647 (PurSolutions) were prepared as described previously (20) and diluted 1:100 in BRB-80 buffer (pH = 7.2) with 20 μM paclitaxel. Diluted MTs were perfused through one channel of the flow cell, incubated for 2 minutes, and then washed with Wash Buffer. This process was then repeated in the same channel, before being repeated twice more in the perpendicular channel. Immediately prior to imaging, both channels were perfused with STORM buffer (Buffer 2 supplemented with 20 μM paclitaxel, 50 mM beta-mercaptoethanol, and 20 mM Cysteamine). After the STORM image was acquired, both flow channels were washed with Motility Buffer. DiO-labelled fluorescent liposomes decorated with an average of 10 kinesins each were then diluted 1:100 in Motility Buffer and perfused into the flow cell and imaged.

Imaging was performed on a Nikon N-STORM microscope equipped with a precise piezoelectric stage and a cylindrical lens as published previously (20, 32). Z-calibration was performed as described previously (40, 41). STORM images of the suspended Alexa-647 MTs were collected by exciting with 405 and 647 nm light and collecting a stack of 20,000 images at a high (∼60 Hz) framerate. Liposomes were imaged using 561 nm excitation at a framerate of 10 Hz. Because the pedestal beads were visible in both the 561 nm and 647 nm channels, they are used as fiducial markers to align the two channels in the XY-plane, while the built in Nikon Perfect Focus system ensured that both image stacks were taken at identical Z heights.

### Data analysis

Data analysis was performed as described previously (20). Briefly, super-resolution particle localizations for the STORM reconstruction of the MTs were performed using the DoM (Detection of Molecules) plugin for ImageJ (42). High resolution tracking of liposomes was also performed using DoM. Liposome localizations were linked into tracks using the “Link Particles to Tracks” function built into DoM. To analyze intersection outcomes and the spatial relationship of each liposome to the intersecting MTs, custom written R scripts were used as described previously (20). Only intersections where the angle between the MTs was between 60° and 120° were included in the analysis. Liposome pausing in 3D intersections was measured by kymography. For a pause to count, motion had to appear stalled in the kymograph for at least 3 frames.

### Size Exclusion Chromatography

200 µg of 1.6 mg/ml mouse kinesin-ΔC with bound kinesin light chain was applied to a 10 x 300 mm Superose 6 (Amersham Biosciences) using an AKTA FPLC (Amersham Pharmacia Biotech). The column was equilibrated with BRB80 (80 mM Pipes pH 7.2, 1 mM EGTA, 1 mM MgCl_2_, 1 mM dithiothreitol) and the protein eluted at a speed of 0.4 ml/min. Each 0.5 ml fraction was concentrated and the total amount in each fraction was applied to a Nupage 4-12% Bis-Tris SDS Gel (Invitrogen). The same experiment was repeated in the presence of 50 µM kinesore (50 mM stock in DMSO) (Tochris Bioscience) in the column equilibration and elution buffers.

### Mechanistic mathematical modeling

The mathematical model and simulation used in this study to predict single and multi-motor transport along single MTs and in 3D MT intersections is identical to that which we previously developed to describe similar transport properties of the shorter, constitutively active kinesin-1 construct, K543 (20). Briefly, the liposome and MTs are modeled as rigid bodies, with each kinesin a 25nm spring-like rod that freely diffuses on the liposome surface and can rotate about its attachment point to the liposome. Kinesin MT attachment occurs at a maximum rate (*k*_*a*,0_) when the distance between the liposome and the MT equals the length of the kinesin and more slowly at shorter and longer distances (20). Diffusion of the kinesins on the liposome surface, and positional and rotational fluctuations of the liposome are modeled explicitly, based on the liposome and kinesin physical properties using an Euler-Maruyama scheme (43–45). Additionally, the force experienced by each kinesin is calculated in each timestep of the simulation, Δ*t* = 5 × 10^−7^*s*. Each kinesin obeys a 3-state mechanochemical kinetic scheme (46), which includes the force dependence of each state transition rate, with chemical reactions simulated at a timestep Δ*t* = 1 × 10^−5^ *s*.

To simulate liposome transport by KinΔC, all parameters in the model were unchanged (20), save either the number of kinesin (N) or the attachment rate (*k*_*a*,0_) as described below to reflect the 3-fold reduction in the attachment rate we observed in the landing assay (see Results and Discussion). Using this model, we simulate unloaded liposome transport along a single MT, transport along a single MT hindered by the load from an optical trap, and liposome transport through a 3D MT-MT intersection. Each simulation begins with a single kinesin bound to the MT and ends when all kinesins detach from the MT, or after 15s has elapsed. We performed two sets of simulations. In the first set, we performed the simulations with N=3 and then again with N=4 (single MT and MT-MT intersection) and with N=7 (optical trap). This approach allowed us to investigate whether our experimental observations with KinΔC could be explained by a 3-fold reduction in motor number with respect to our observations with K543, since we can explain these latter observations with the model having N=10 kinesin motors in the single MT and MT intersection measurements, and N=20 kinesin motors in the laser trap (20). In the second set of simulations, we performed simulations with a 3-fold reduction in attachment rate, *k*_*a*,0_ = 50s^-1^, compared to the model that successfully described our observations with K543 which used *k*_*a*,0_ = 150s^-1^.

## Results and Discussion

### KinΔC-liposomes go straight or terminate but rarely turn in 3D MT intersections

Kinesin-1 motors must navigate their vesicular cargo through multiple 3D MT intersections en route to the cargo’s destination (21–24). If kinesin-1 motors exist in an equilibrium between the autoinhibited and active states (Fig. 1A) while attached to a vesicle, does this equilibrium impact how the vesicle is maneuvered through a MT intersection? To address this question *in vitro,* we suspended MTs between poly-L-lysine-coated silica beads on a coverslip surface, creating orthogonal 3D MT intersections (Fig. 1B and C). We then decorated 350-nm fluid-like (DOPC) liposomes (Fig. 1B) with one of two kinesin-1 constructs that could freely diffuse on the liposome surface, which were present at ∼10 motors per liposome, based on a photobleaching-based counting assay (Fig. S1). The ubiquitously expressed murine kinesin-1 heavy chain, KIF5B, was the basis for both constructs (Fig. 1A). The KinΔC construct (1-888 amino acids) with full-length KLC2 bound, retains the capacity to be partially autoinhibited, based on a 3-fold lower landing rate compared with the constitutively active truncated construct K543 (Fig. 3C). Both constructs are C-terminally biotinylated to provide an attachment handle to the liposome (20, 25).

**Figure 1.**
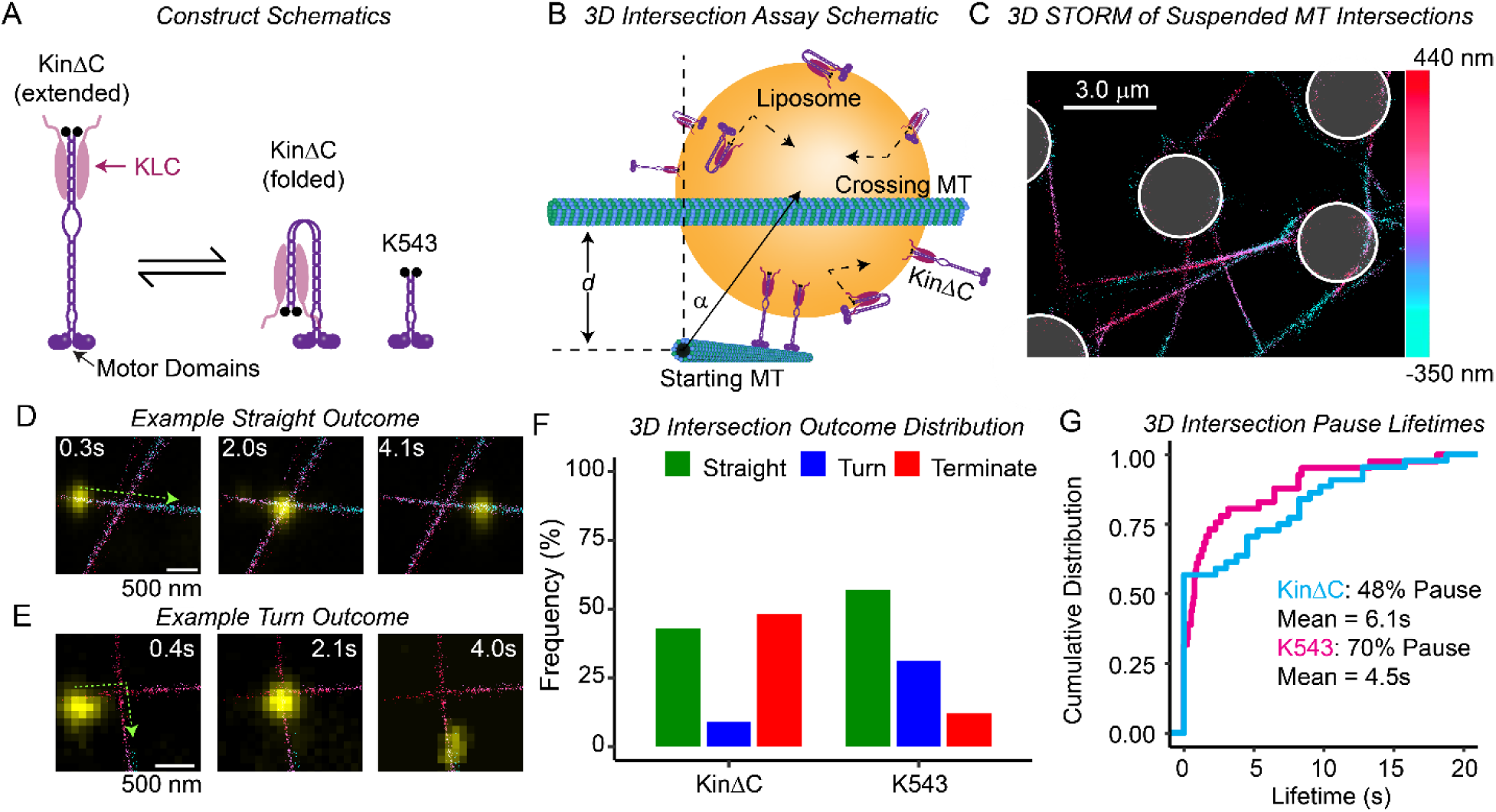
*KinΔC terminates frequently in 3D MT intersections.* (A) Schematic of constructs of interest. KinΔC (left) contains aa1-888 of murine KIF5B and copurifies with full-length KLC2. Critically, KinΔC is able to adopt the folded conformation associated with autoinhibition. K543 contains aa1-543 of murine KIF5B and serves as a constitutively active control. Both constructs are C-terminally biotinylated (black circles). (B) Schematic cartoon of 3D MT intersection assay. Liposome (gold) is transported by KinΔC (purple) along MTs (green) while KinΔC motors can diffuse on the liposome surface. The liposome is moving out of the page and approaches the horizontal MT in a 3D intersection. The geometry of the interaction between the liposome and the intersection is described by two parameters, *d* and α, illustrated here, where *d* is the vertical gap between the intersecting MTs, and α is the angle of approach of the liposome coming in to the intersection, where an α of 0° indicates the liposome is pointed up towards the crossing MT, and an α of 180° indicates the liposome is pointed down and away from the crossing MT. (C) 3D STORM reconstruction of suspended MT intersections. Z-position of MTs is shown via the color bar on the right. (D) Example of a liposome (yellow) passing straight through a 3D intersection. Color bar as in C. Gap = 34 nm. (E) Example of a liposome (yellow) turning in a 3D intersection. Color bar as in C. Gap = 174 nm. (F) Bar graph of 3D intersection outcomes for liposomes with 10 KinΔC (left) compared to 10 K543 control (right). 10 K543 control data are replotted from Bensel et al., 2024. For KinΔC data, N = 57 intersection outcomes from 3 independently performed intersection assays. (G) Cumulative distribution plot of the pause lifetimes observed for liposomes with 10 KinΔC (cyan) and 10 K543 (magenta) in 3D intersections. Events with no discernible pause are represented as a pause lifetime of 0s. K543 lifetime data are replotted from Bensel et al., 2024.

**Figure 2.**
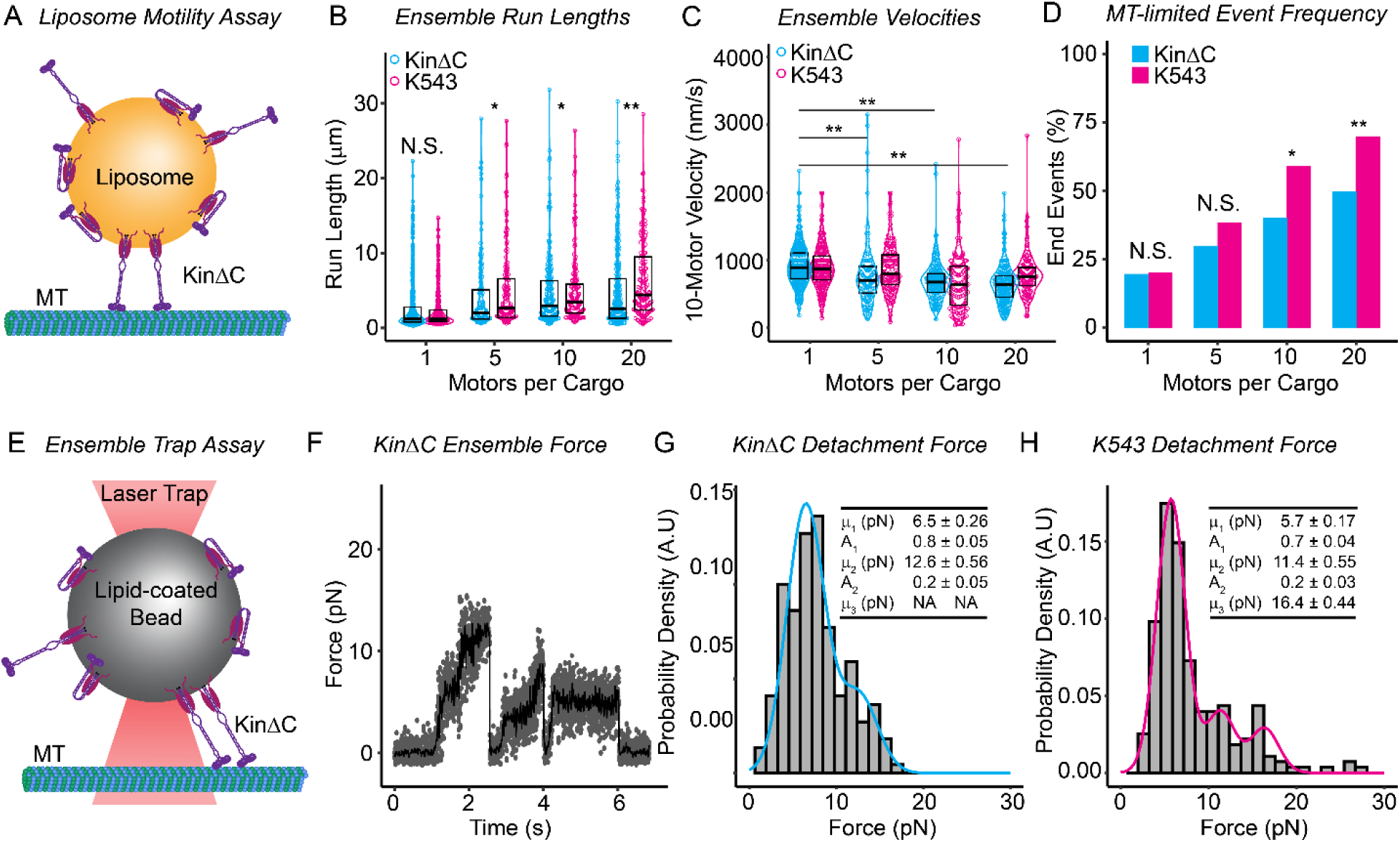
*KinΔC Ensembles Favor Fewer Engaged Kinesins than K543 Ensembles.* (A) Schematic representation of multi-motor motility assay with fluid-like liposome as cargo. Liposome (gold) has 5-20 kinesins (purple) bound to its surface. The fluorescence of the liposome is tracked by TIRF microscopy as it is moved along MTs (green) by the bound kinesins. (B) Dot plots with overlaid violin plots representing multi-motor run lengths measured for KinΔC (cyan) and K543 (magenta) with up to 20 bound kinesins. Black overlaid box plot shows median and upper and lower quartiles per condition. K543 control data are replotted from Bensel et al., 2024. p-Values are as follows: N.S.: p>0.05, *: p<0.05, **: p<0.01. Run length data are compared using a Kruskal-Wallis with Dunn’s post-hoc test using the Benjamin-Yuketieli adjustment. See Table S1 for exact p-values. For KinΔC data, N_events_ ranges from 137 – 480 from at least 3 independent experiments. (C) Dot plot with overlaid violin plot representing multi-motor velocities measured for KinΔC (cyan) and K543 (magenta). Black overlaid box plot shows median and upper and lower quartiles per condition. K543 control data are replotted from Bensel et al., 2024. p-Values are as follows: N.S.: p>0.05, *: p<0.05, **: p<0.01. Velocity data are compared using a Kruskal-Wallis with Dunn’s post-hoc test using the Benjamin-Yuketieli adjustment. See Table S1 for exact p-values. N’s are the same as in (B). (D) Bar graph depicting the percentage of runs for each condition (single kinesin Qdot, and 5, 10, and 20 kinesins per liposome) which reach the end of the MT before detaching. K543 data are replotted from Bensel et al., 2024. End event frequencies are compared using a two proportions Z-test. p-Values are as follows: N.S.: p>0.05, *: p<0.05, **: p<0.01. See Table S1 for exact p-values. N’s are the same as in (B, C). (E) Schematic representation of multi-motor optical trapping assay with lipid-coated bead as cargo. Kinesins (purple) are bound to lipid-coated beads (grey) in a 20 to 1 excess. The laser trap (red) is used to position the bead close to a MT (green) and force ramps are recorded. (F) Sample force ramp collected for a lipid-coated bead coated with 20 KinΔC motors. Raw data is shown in grey with median-filtered data overlaid in black. (G) Detachment force histogram with overlaid double Gaussian fit collected for KinΔC experimental results (cyan). Fit parameters ± fitting error are shown in inset. N_events_ = 169 from 5 independent experiments. (H) Detachment force histogram with overlaid triple Gaussian fit collected for K543 (magenta). K543 data and fit are replotted from Bensel et al., 2024. Fit parameters ± fitting error are shown in inset.

**Figure 3.**
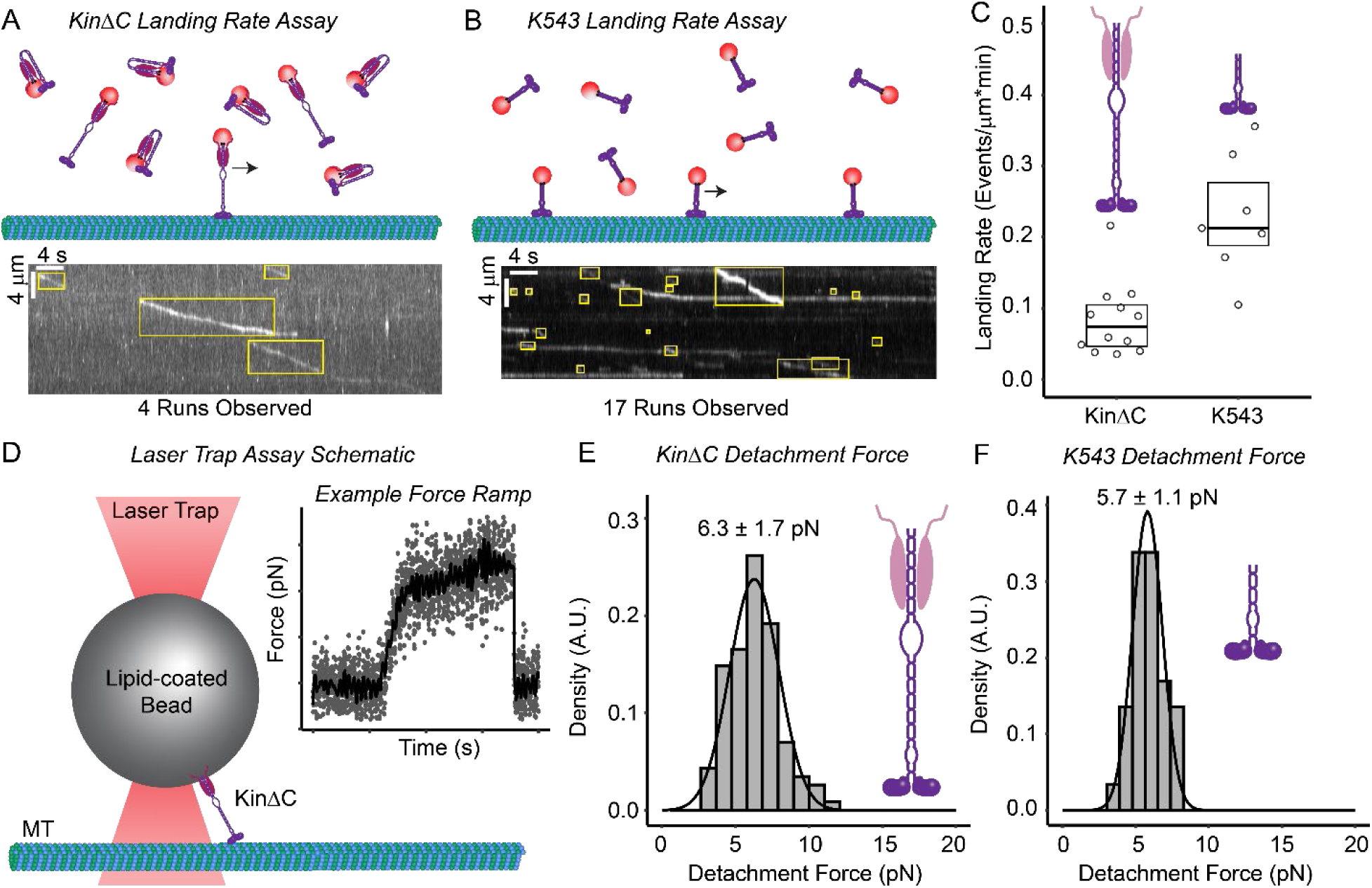
*KinΔC and K543 are indistinguishable once engaged with the MT.* (A, upper) Schematic of landing rate assay as performed with KinΔC. KinΔC is bound to Qdots (red) and flowed into a microscopy chamber containing MTs (green). (A, lower) Example kymograph of single molecule Qdot assay with KinΔC. Runs are boxed in yellow. Distance along MT is represented by the vertical axis while time is represented as the horizontal axis. A moving particle appears as a slanted line while a stuck particle appears as a vertical line. (B, upper) Schematic of landing rate assay as performed with K543. K543 is bound to Qdots (red) and flowed into a microscopy chamber containing MTs (green). (B, lower) Example kymograph of single molecule Qdot assay with K543. Runs are boxed in yellow. Axes are as in (A). (C) Dot plot with overlaid box plot for landing rate determined for KinΔC (left) and K543 (right). Each dot represents the landing rate determined for a single MT in a single video with the equation: Landing Rate = N_runs_/[MT length (μm) x video length (min)]. Statistical analysis was performed using Kruskal-Wallis with Dunn’s post-hoc test using Benjamin-Yuketieli adjustment, p = 0.01. N= 12 MTs (KinΔC) or 7 MTs (K543) from 3 independent videos each. (D) Schematic representation of single-molecule optical trapping assay as performed. A lipid-coated silica bead (grey) with a bound KinΔC (purple) is brought close to a surface-bound MT (green) by the laser trap (red). (D, Inset) Sample force ramp collected from a single kinesin-ΔC molecule in the optical trapping assay. Median-filtered trace (black) is overlaid on top of raw data (grey). (E) Histogram with overlayed fit of detachment forces measured for single KinΔC molecules. KinΔC has a fitted mean detachment force of 6.3 ± 1.7 pN. N_events_ = 109 from 4 independent experiments. (F) Histogram with overlayed fit of detachment forces measured for single K543 molecules. K543 has a fitted mean detachment force of 5.7 ± 1.1 pN. K543 run detachment force data are replotted from Bensel et al., 2024.

Using 3D stochastic optical reconstruction microscopy (STORM), the liposome’s spatial relationship to the MT intersection was defined, i.e., the liposome approach angle (α) and vertical gap (*d*) between the MTs (Fig. 1B and C) (20, 30). This ensured that we only considered directional outcomes (i.e., whether the liposome went straight (Fig. 1D, Video S1), turned (Fig. 1E, Video S2), or terminated its run) that resulted from motors on the liposome surface being able to reach the intersecting MT. For each event that met our criteria, we recorded if the liposome paused, how long it paused, and the directional outcome.

At 3D MT intersections, KinΔC-liposomes were nearly as likely to terminate (48%) as to go straight (43%) but rarely turned onto the crossing MT (9%) (Fig. 1F). These directional outcomes were distinctly different than we previously reported for K543-liposomes, which preferred to go straight (58%) or turned occasionally onto the crossing MT (37%), but rarely terminated (5%) (Fig. 1F) (20). Could pausing in the intersection offer insight to these constructs’ different directional outcomes? In fact, KinΔC-liposomes were less likely to pause (48% paused) compared to K543-liposomes (70% paused) (20), but when paused their lifetimes (6.1 sec) were 36% longer (Fig. 1G). Our previous *in silico* modeling of K543 transport in 3D MT intersections suggested that pausing originates from motors on the liposome surface binding simultaneously to both MTs and engaging in a tug-of-war, which must be resolved before a directional outcome is determined (20, 23, 43). The observation that KinΔC-liposomes rarely turn suggests that the motors’ autoinhibition reduces the number of available KinΔC motors that can engage the crossing MT resulting in less frequent tug-of-wars, pausing, and turning events. If resolving a tug-of-war requires MT engagement of a new motor, a reduction in the number of available motors would also prolong these events resulting in longer pauses.

### KinΔC ensembles have reduced MT engagement

For kinesin-1 transported cargoes, the run length, or characteristic travel distance, scales with the number of motors that are MT-engaged (47–50), while velocity slows due to mechanical coupling creating internal loads between the motors (20, 51, 52). Therefore, we measured KinΔC-liposome run lengths and velocities along single MTs adhered to a glass slide for varying size KinΔC ensembles (∼5, ∼10, or ∼20 motors, Fig. 2A) compared to a single KinΔC motor transporting a quantum dot (Qdot) cargo. As expected, KinΔC ensembles universally had longer run lengths (1.99 – 2.98 μm) and slower velocities (637 – 702 nm/s) than single KinΔC motors (1.19 μm and 890 nm/s) (Fig. 2B and C). However, compared to K543-liposomes transported by similar ensemble sizes, KinΔC-liposomes had shorter median run lengths (Fig. 2B, Table S1), although velocities were similar (Fig. 2C, Table S1). In fact, these run length differences may be underestimated, as run lengths can be limited by the MT length itself (34–36). Knowing that MTs were prepared identically with similar length distributions (Fig. S2), we compared the frequency at which liposomes reached the MT end (Fig. 2D). For all motor ensemble sizes, K543-liposomes were ∼46% more likely than KinΔC-liposomes to reach the MT end (Fig. 2D), suggesting that the K543-liposomes run lengths are more highly underestimated than those of the KinΔC-liposomes. Therefore, the reduction in KinΔC-liposome run lengths compared to K543-liposomes is even greater. These data support the hypothesis that fewer KinΔC motors from the ensemble engage the MT compared to the constitutively active K543 control.

To effectively count how many KinΔC motors in the ensemble are MT engaged, we measured the cargo’s detachment force in an optical trapping assay, as this force should scale with the engaged motor number (48, 49, 53) (Fig. 2E). Therefore, we incubated 500-nm lipid-coated beads with 20-fold excess KinΔC to match the motor density of a 350-nm liposome with 10 bound motors and then measured the detachment force as the KinΔC motors step against the hindering load of the trap (Fig. 2E and F). The KinΔC-bead detachment forces were distributed as the sum of two Gaussians centered at 6.5 and 12.6 pN, with 80% of the events in the first peak (Fig. 2G), whereas K543-bead detachment forces were described by a triple Gaussian with peaks at 5.7, 11.4, and 16.4 pN (Fig. 2H). The initial detachment force peaks for the KinΔC and K543 constructs were equivalent to that measured for a single motor (KinΔC: 6.4 ± 1.9 pN; K543: 5.7 ± 1.1 pN) (Fig. 3E and F) and similar to that in the literature (54–56). Therefore, often only 1 but at most 2 KinΔC motors are MT engaged compared to K543 ensembles where up to 3 motors can be MT engaged. These data once again suggest that autoinhibition effectively reduces the number of active motors.

### KinΔC has a reduced MT landing rate but once MT engaged is indistinguishable from K543

If kinesin-1 can transition between autoinhibited and active states, does this dynamic equilibrium affect how often the motor engages a MT and its motile properties once engaged with the MT? To quantify MT engagement, we measured the landing rate (number of processive runs per unit time per unit MT length) of single Qdot-labelled KinΔC (1 nM) relative to K543 (1 nM) in saturating MgATP (Fig. 3A and C). KinΔC initiated processive runs at a rate (0.08 μm^-1^ min^-1^) ∼3-fold less than K543 (0.23 μm^-1^ min^-1^) (Fig. 3B and C, Video S3), suggesting that kinesin-1 autoinhibition reduces MT engagement. Once engaged, the mechanochemical properties of single KinΔC motors were not impacted by autoinhibition, with median run lengths (RL), mean velocities (V) and detachment forces (F) (Fig. 3D) (RL= 1.19 μm; V=890 nm/s; F= 6.4 ± 1.9 pN) (Fig. 2B and C, Fig. 3E) that were indistinguishable from K543 (RL= 1.19 μm; V= 868 nm/s; F= 5.7 ± 1.1 pN) (Fig. 2B and C, Fig. 3F). Once KinΔC engages with the MT, the motor’s processivity and force generation are governed by the mechanochemistry of the fully active motor, suggesting that transitions back into the autoinhibited state must occur only in solution, following MT detachment. If the reduced rate of MT association reflects a dynamic equilibrium between the autoinhibited and active states, these data do not inform how rapidly the two states exchange (see modeling section below). Nonetheless, the reduced landing rate of KinΔC suggests that autoinhibition can effectively reduce the number of MT engaged motors in an ensemble and thus the ensemble’s cargo transport capacity as described above.

### Kinesore treatment activates an autoinhibited KinΔC motor

If autoinhibitory intramolecular interactions are responsible for KinΔC’s reduced MT engagement, disrupting these interactions should activate the motor. To test this idea we used kinesore, a small molecule that interacts with the KLC to mimic cargo binding and disrupt inhibitory intramolecular interactions to activate kinesin-1 (26, 57). In the presence of 50 μM kinesore, KinΔC’s landing rate increased significantly to 0.26 μm^-1^ min^-1^, equal to that of K543 (0.22 μm^-1^ min^-1^) (Fig. 4A and B, Video S3 and S4), which was unaffected by kinesore as this construct lacks KLC. By restoring the KinΔC MT landing rate, would kinesore treatment of a KinΔC ensemble increase its force-bearing capacity (i.e., increased number of MT-engaged motors)? This was in fact the case as the distribution of detachment forces measured for KinΔC-beads in the optical trap with 50 μM kinesore present was now best fit by the sum of three rather than two Gaussians, centered at 7.2 pN, 13.7 pN, and 20.0 pN (Fig. 4C and D), similar to that for K543-beads (Fig. 2H). Therefore, the number of MT engaged KinΔC motors increased from at most 2 in the absence of kinesore to up to 3 engaged motors with kinesore present.

**Figure 4.**
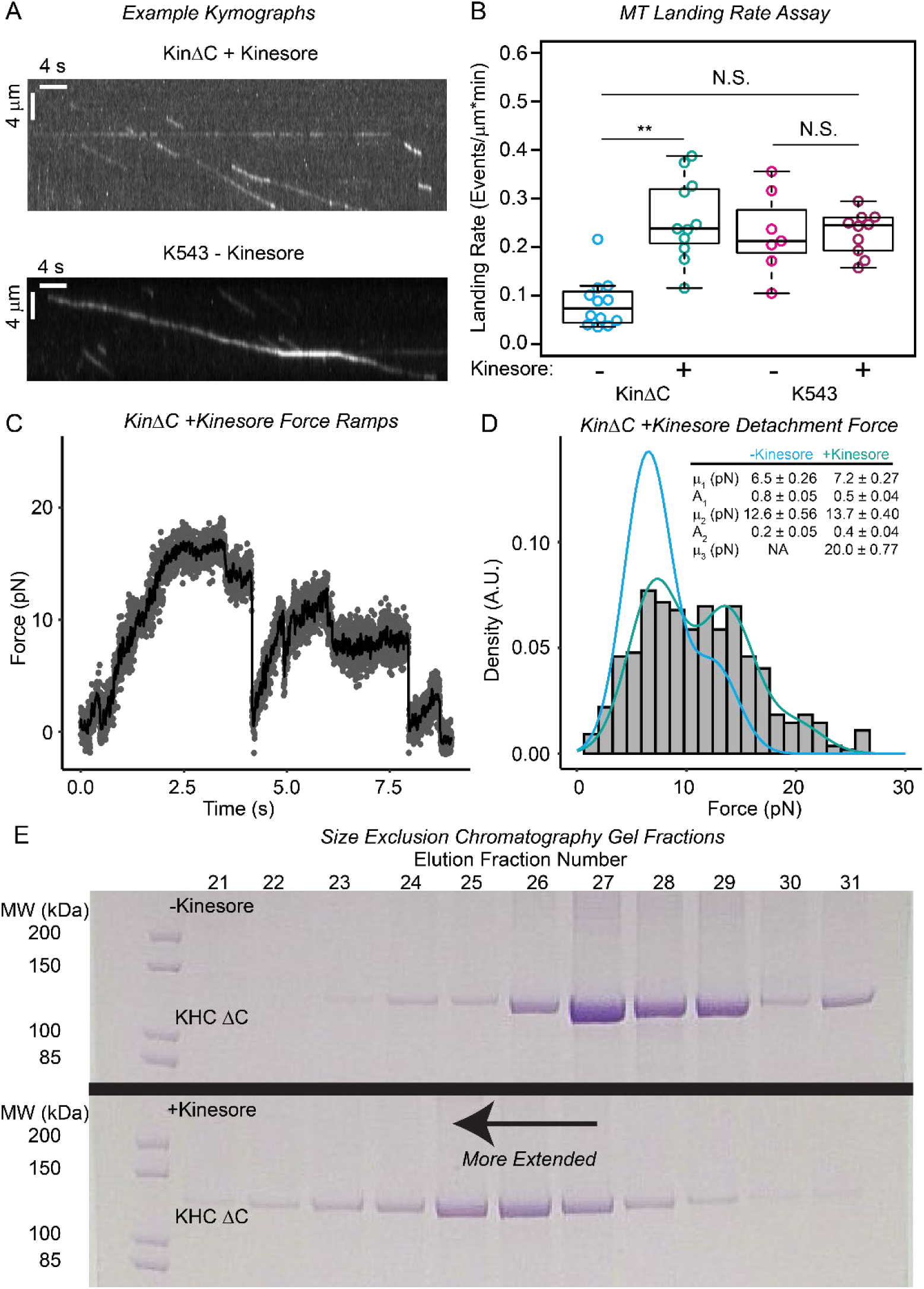
*Kinesore restores the MT engagement activity of KinΔC to that of K543.* (A) Representative landing rate assay kymographs for KinΔC (top) and K543 (bottom) in the presence (top) or absence (bottom) of 50 μM kinesore. Distance is shown on the horizontal axis, while time is shown on the vertical axis. (B) Dot plot of landing rates measured in the absence (-) and presence (+) of 50 μM kinesore for KinΔC (cyan, turquoise) and K543 (magenta, burgundy). Each dot represents the landing rate determined from a single MT in a single video. Statistical analysis of landing rate data was performed using Kruskal-Wallis with Dunn’s post-hoc test using the Benjamin-Yuketieli adjustment. p-Values are as follows; N.S.: p>0.05, **: p<0.01. KinΔC + kinesore, N_events_ = 11 MTs from 3 independent videos, K543 + kinesore, N_events_ = 10 MTs from 3 independent videos. (C) Sample force ramp recorded from a lipid-coated bead being moved by 20 KinΔC motors in the presence of kinesore. Raw data are shown in grey and median filtered data are overlaid in black. (D) Histogram with overlaid triple Gaussian fit for detachment forces measured KinΔC in the presence of kinesore (turquoise) with the double Gaussian fit to detachment forces measured for KinΔC in the absence of kinesore (cyan) shown as reference. N_events_ = 418 from 4 independent experiments. (E) SDS-PAGE of size exclusion chromatography elution fractions for KinΔC in the absence (top) or presence (bottom) of 50 μM kinesore. Elution fractions are shown increasing from left to right, the band corresponding to the KinΔC heavy chain (KHC ΔC) is shown. Given the same molecular weight, a molecule will elute sooner the more extended it is.

Implicit in the activation of KinΔC by kinesore is that disrupting the autoinhibitory intramolecular interactions should result in a more extended molecule (15, 17, 26). We utilized size exclusion chromatography to assess the relative shape of KinΔC molecules in the presence or absence of 50 μM kinesore. A molecule with an extended structure elutes sooner than a molecule of the same molecular mass with a folded structure (Fig. 4E). Elution fractions from the size exclusion chromatography column equilibrated and eluted in the absence (Fig. 4E, top) versus presence of 50 μM kinesore (Fig. 4E, bottom) show that KinΔC elutes earlier in the presence of kinesore. These data suggest that kinesore binding to KLC results in a more extended conformation, consistent with an active KinΔC motor.

### KinΔC autoinhibition: mechanistic model and impact on liposome transport

The transport data for single KinΔC motors and ensembles together suggest that KinΔC can adopt the folded and extended states even when attached to cargo. To understand how KinΔC autoinhibition impacts ensemble transport, we took advantage of our previously published mechanistic *in silico* model that predicted single motor and ensemble K543 transport (Fig. 5A) on single MTs and in 3D MT intersections (see Methods for details of simulations) (20). Crucially, the model is predictive because it is constrained only by a three-state model of kinesin’s mechanochemistry (46) and the physical properties of the liposomes and MTs. Adopting this model to predict KinΔC transport was simplified knowing that once engaged with the MT, the mechanochemical properties of KinΔC are identical to K543. Therefore, the *in silico* model was modified to include KinΔC autoinhibition as a dynamic equilibrium between an inhibited and an active state where the exchange rates between states are either slow on the timescale of the simulation such that KinΔC exists predominantly in the inhibited state (i.e. effectively reducing the number of active motors in the ensemble) (Fig. 5E) or the exchange rates are much faster than the timescale of the simulation such that autoinhibition can be modeled simply as a reduced apparent attachment rate (Fig. 5F). In support of these two concepts, others in the field have proposed both motor number (58) and reattachment kinetics (50, 59) as important determinants of ensemble transport properties.

**Figure 5.**
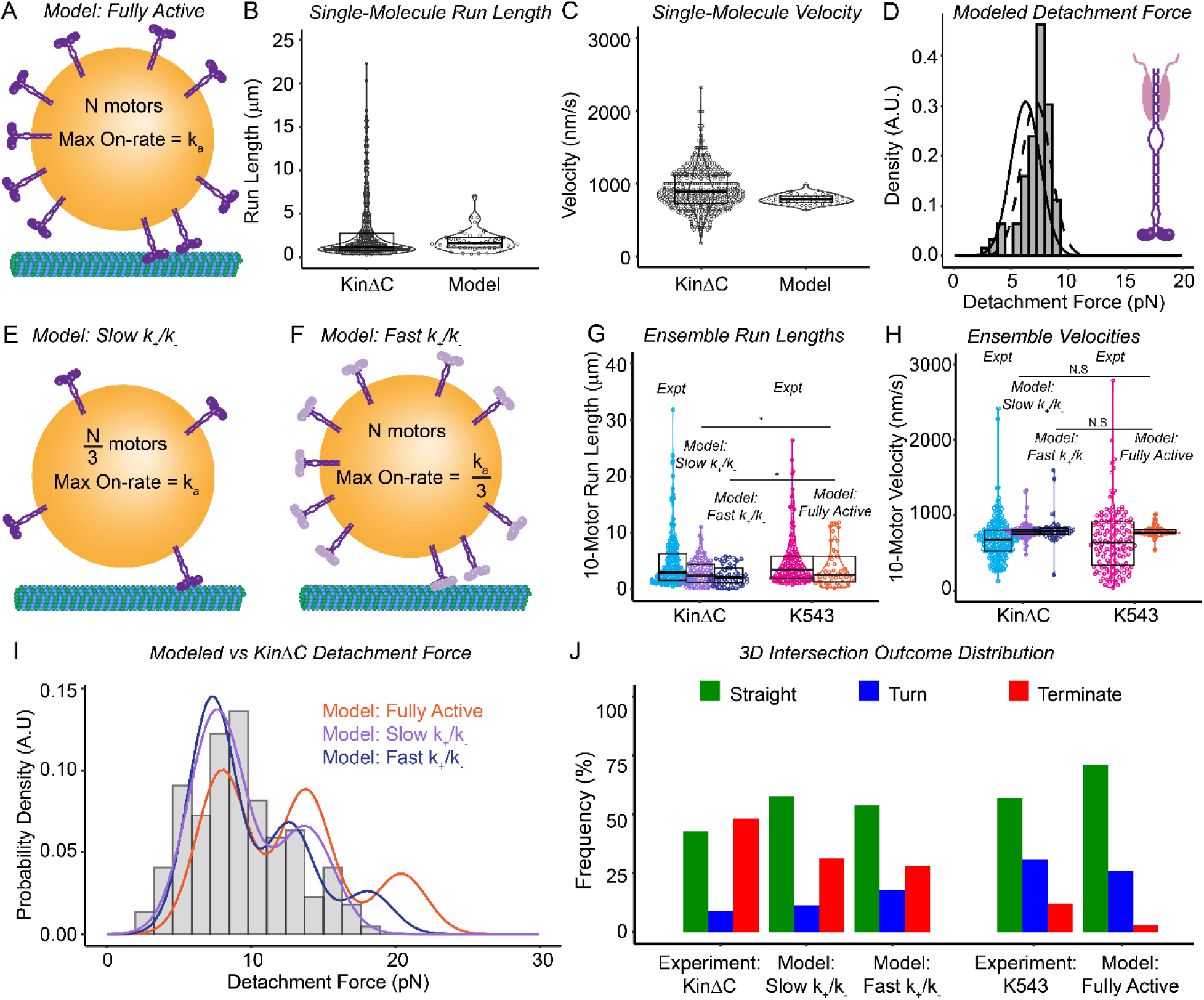
*Mechanistic modeling of kinesin-1 autoinhibition predicts experimental differences observed between KinΔC and K543.* (A) Schematic representation of our mechanistic model of kinesin-1 transport as published in Bensel et al., 2024. All motors in the model are active and bind MTs with a maximum rate k_a_ = 150 s^-1^. We refer to this as the ‘fully active’ model. (B) Dot plot with overlaid box plot and violin plot of KinΔC (left) and modeled (right) single molecule run length. Model N_events_ = 46. (C) Dot plot with overlaid box plot and violin plot of KinΔC (left) and modeled (right) single molecule velocity. Model N_events_ = 46. (D) Histogram of modeled detachment force distribution with overlaid single Gaussian fit (black, dashed). Single Gaussian fit to KinΔC detachment force experimental data shown for reference (black, solid). Model N_events_ = 99. (E) Schematic representation of our modified mechanistic model of kinesin-1 which emulates an autoinhibited motor where exchange between the active and inhibited states occurs slowly. We call this model “Slow k_+_/k_-_“ and it is implemented by reducing the number of motors in the simulation by a factor of 3, i.e., N/3 motors. All other model parameters remain the same. (F) Schematic representation of our modified mechanistic model of kinesin-1 which is modified to capture an autoinhibited motor that exchanges rapidly between the active and inhibited states. We call this model “Fast k_+_/k_-_“ and it is implemented by reducing maximum rate of MT association by a factor of 3 to k_a_ = 50 s^-1^, but no other parameters of the model are changed. (G) Dot plot with overlaid violin plot and box plot of run lengths measured for experimental (Expt) and modeled liposomes with ∼10 bound motors. KinΔC experimental results are shown for reference (cyan) next to the model of Slow k_+_/k_-_ (purple) (N_events_ = 93) and Fast k_+_/k_-_ (dark blue) (N_events_ = 50). K543 experimental results (magenta) and fully active model results (orange) are replotted from Bensel et al., 2024 and shown for reference. Statistical analysis of run length data was performed using Kruskal-Wallis with Dunn’s post-hoc test using the Benjamin-Yuketieli adjustment. p-Values are as follows; N.S.: p>0.05, *: p<0.05. (H) Dot plot with overlaid violin plot and box plot of velocities measured for experimental (Expt) and modeled liposomes with ∼10 bound motors. KinΔC experimental results are shown for reference (cyan) next to the model of Slow k_+_/k_-_ (purple) and Fast k_+_/k_-_ (dark blue). K543 experimental results (magenta) and fully active model results (orange) are replotted from Bensel et al., 2024 and shown for reference. Statistical analysis of run length data was performed using Kruskal-Wallis with Dunn’s post-hoc test using the Benjamin-Yuketieli adjustment. p-Values are as follows; N.S.: p>0.05. (I) Histogram of experimentally measured detachment forces for ensembles of 20 KinΔC with overlaid model prediction fits. The model of Slow k_+_/k_-_ is shown in purple (N_events_ = 200), Fast k_+_/k_-_ is shown in dark blue (N_events_ = 200), and fully active model is shown in orange. Fully active model fit is replotted from Bensel et al., 2024. (J) Bar graph of 3D intersection outcomes for experiments and simulations. Slow k_+_/k_-_ N_events_ = 285, fast k_+_/k_-_ N_events_ = 260. K543 experimental results and fully active model results are replotted from Bensel et al., 2024.

For single KinΔC motors, the model predicts run lengths, velocity, and detachment forces (RL= 1.66 μm; V= 787 nm/s; F= 6.5 pN) as observed experimentally and as reported previously for K543 (20) (Fig. 5B-D). To model autoinhibition and its effect on KinΔC MT engagement, we assumed that the 3-fold lower KinΔC MT landing rate compared to K543 (Fig. 3C) was due to autoinhibition, as supported by the observation that kinesore restores the landing rate of KinΔC to that of K543 (Fig. 4B). Therefore, we first ran liposome transport simulations, as described above, with autoinhibition reducing the number of active motors in the ensemble by a factor of 3 to model a slow exchange rate between the active and inhibited states (Fig. 5E). Since liposomes were decorated with a 10-motor ensemble, we simulated liposome transport by either 3 or 4 motors and combined the results for each transport condition to best reflect the desired 3.33 motors (i.e. 10 motors divided by a factor of 3). Alternatively, we set the maximum MT attachment rate (*k*_*a*,0_ = 50s^-1^) for all motors in the ensemble to 1/3 of the rate which predicted the K543 observations (*k*_*a*,0_ = 150s^-1^), reflecting a 3-fold reduction in the apparent attachment rate and modeling rapid exchange between the active and inhibited states (Fig. 5F).

Simulations for reduced motor number and MT attachment rate reproduced the experimental trends in ensemble run lengths and velocities for KinΔC versus K543 (Fig. 5G and H). Interestingly, the reduced motor number simulations were consistent with the experimentally observed detachment force distributions showing most often 1 or at most 2 MT engaged KinΔC motors (Fig. 5I). Simulations with a reduced MT attachment rate, although broadly similar to the reduced motor number case, did have a minor population of 3 simultaneously engaged motors, though less frequently than in the active model (Fig. 5I). For KinΔC-liposomes in 3D MT intersections, both the reduced motor number and MT attachment rate simulations qualitatively reproduced the predominant directional outcomes of going straight through or terminating in the intersection that were dramatically different than the K543-liposome directional outcomes and those predicted by a model of fully active motors (Fig. 5J) (20).

To understand why reduced motor number or MT association rate affects directional outcomes in 3D MT intersections, we must rely on our *in silico* model to provide a mechanistic basis for how each directional outcome occurs. In brief, as a liposome enters the intersection, motors on the liposome surface can bind simultaneously to both MTs and thus the liposome may pause if the motors engage in a tug-of-war. In a tug-of-war, motors stochastically attach and detach from both MTs with the tug-of-war resolved when motors on one of the intersecting MTs all detach, allowing the winning team to pull the liposome straight (Video S5) or turn (Video S7). Generally, if no tug-of-war occurs, the liposome proceeds straight (Video S6). Finally, if all MT engaged motors detach, transport is terminated (Video S8). If we reduce motor-MT engagement by either reducing the number of available motors or the MT attachment rate, a tug-of-war is less likely to occur, favoring straight outcomes. Furthermore, since both experimental and simulated data suggest that KinΔC-liposome transport is driven most often by a single engaged motor, when the liposome collides with the crossing MT the motor may detach, terminating the run (Video S9). Thus, our model provides a mechanistic basis for kinesin autoinhibition impacting directional outcomes at 3D MT intersections. Interestingly, modelling autoinhibition as an equilibrium between a folded and extended state where the exchange rates between the states are either slow or rapid (see above) captures much of the KinΔC transport behavior (Fig. 5), suggesting that this mode of motor regulation is not sensitive to the exchange rates. However, the detachment force distribution is better predicted by the model of slow exchange rates suggesting that the folded, autoinhibited state resulting from multiple intramolecular interactions is most likely quite stable, as reported in the literature (15–18).

## Conclusion

Here we provide evidence that kinesin-1 autoinhibition has profound effects on liposome transport *in vitro* whether it be on a single MT or when challenged by a 3D MT intersection. The partial autoinhibition exhibited by the KinΔC construct is only one example of what is likely a range of motor activation that is possible in cells and depends on kinesin-1 isoform (16), whether or not KLCs are bound (16, 17), the cargo adapter that kinesin is bound to (16, 19), as well as MAPs such as MAP7 that enhance engagement of kinesin with the MT (16, 60, 61). While this study focuses on kinesin-1, the mechanochemical differences in kinesin subfamilies may provide the cell additional capacity to tune cargo transport (59, 62, 63). Given the cell’s 3D MT network with numerous intersections, optimizing the directional outcome at each intersection, which is sensitive to the effective number of active motors, provides the cell a mechanism for ensuring vesicular cargo delivery to the proper destination.

## Data and code availability

Due to the size of the super resolution imaging datasets, data are stored on University of Vermont servers. The data are available from the corresponding author upon reasonable request. Modeling code and analysis scripts can be made available from the corresponding author upon reasonable request.

## Author contributions

B.M.B, K.M.T., S.W., and D.M.W designed research; B.M.B., S.B.P., P.M.F., and S.W. performed research, B.M.B., S.W., K.M.T., and D.M.W. analyzed data, and B.M.B., K.M.T., S.W., and D.M.W. wrote the paper.

## Acknowledgements

We would like to acknowledge Shane R. Nelson for supporting in developing data analysis scripts, M. Yusuf Ali for experimental input, Guy Kennedy for TIRF microscopy support and training, Andrew Lombardo for guidance on experiments and data analysis, Douglas Taatjes and Nicole Bouffard of the UVM Microscopy Imaging Center (RRED# SRC_018821) for training and support in 3D N-STORM imaging and analysis. We would also like to thank current and former members of the Warshaw and Trybus labs for their invaluable input, discussions, and support. The work published here would not have been possible without the contributions of those who have played a role in the creation, distribution, and maintenance of the open-source software packages used in this study, particularly r-project.org and ImageJ.nih.gov. We also thank our funding: NIH Grant T32HL076122 (to B.M.B.), NIH Grant F32GM140618 (to B.M.B.), NIH Grant R35GM141743 (to D.M.W.), NIH Grant R35GM136288 (to K.M.T.), and NIH Grant S10OD026884 (to D.M.W.), and a generous gift from Arnold and Mariel Goran to D.M.W.

## Abbreviations

2D: 2-dimensional
3D: 3-dimensional
K543: kinesin 543
KinΔC: kinesin-ΔC
KHC: Kinesin heavy chain
KLC: Kinesin light chain
MT: Microtubule
Qdot: Quantum dot
ROI: Region of interest
STORM: Stochastic optical reconstruction microscopy

## Supplemental Movies

Movie S1. *KinΔC liposomes can go straight in 3D MT intersections.* Liposome (yellow) with ∼10 bound KinΔC passes straight through a suspended 3D MT intersection. The liposome channel is overlaid with the 3D STORM reconstruction of the MTs where the color of the MTs gives the position of that point in Z according to the color bar shown in Fig. 1C. Scale bar: 500 nm; collected at 10 Hz; playback at 1X real time.

Movie S2. *KinΔC liposomes can turn in 3D MT intersections.* Liposome (yellow) with ∼10 bound KinΔC turns in a suspended 3D MT intersection. The liposome channel is overlaid with the 3D STORM reconstruction of the MTs where the color of the MTs gives the position of that point in Z according to the color bar shown in Fig. 1C. Scale bar: 500 nm; collected at 10 Hz; playback at 1X real time.

Movie S3. *KinΔC lands on MTs at a low rate compared to K543.* Video of KinΔC (left) and K543 (right) single molecule motility assays. Fluorescent MTs are shown in cyan while Qdot fluorescence is shown in red. Scale bar: 5 mm; collected at 10 Hz; playback at 10X real time.

Movie S4. *Kinesore elevates the landing rate of KinΔC.* Video of KinΔC single molecule motility assay with the addition of 50 mM kinesore. Fluorescent MTs are shown in cyan while qdot fluorescence is shown in red. Scale bar: 5 mm; collected at 10 Hz; playback at 10X real time.

Movie S5. *Simulated liposomes with reduced motor number can go straight in 3D intersections with a tug-of-war.* Video of simulated liposome with 4 liposome-bound kinesins going straight in a 3D MT intersection. The liposome is shown in gold with the blue band indicating the hemisphere of the liposome. MTs are shown in green. Kinesins attached to the liposome but not engaged with the MT are shown in magenta. Kinesins engaged with the MT are shown as red and yellow heads with a magenta stalk. A red line represents the attachment between the kinesin tail and the liposome and the length of the line represents the force on that attachment point. Notice that the liposome avoids the crossing MT because motors engaged with the crossing MT pull the liposome around it, then detach, allowing the motors engaged with the starting MT to pull the liposome straight through the intersection.

Movie S6. *Simulated liposomes with reduced motor number can go straight in 3D intersections without a tug-of-war.* Video of simulated liposome with 3 liposome-bound kinesins going straight in a 3D MT intersection. The liposome is shown in gold with the blue band indicating the hemisphere of the liposome. MTs are shown in green. Kinesins attached to the liposome but not engaged with the MT are shown in magenta. Kinesins engaged with the MT are shown as red and yellow heads with a magenta stalk. A red line represents the attachment between the kinesin tail and the liposome and the length of the line represents the force on that attachment point. Notice that the liposome avoids the crossing MT by simply wobbling around it.

Movie S7. *Turning in 3D intersections requires a tug-of-war.* Video of simulated liposome with 3 liposome-bound kinesins turning in a 3D MT intersection after the kinesins engaged with the intersecting MT win the tug-of-war. The liposome is shown in gold with the blue band indicating the hemisphere of the liposome. MTs are shown in green. Kinesins attached to the liposome but not engaged with the MT are shown in magenta. Kinesins engaged with the MT are shown as red and yellow heads with a magenta stalk. A red line represents the attachment between the kinesin tail and the liposome and the length of the line represents the force on that attachment point.

Movie S8. *Simulated liposomes with reduced motor number can terminate in intersections with a tug-of-war.* The liposome is shown in gold with the blue band indicating the hemisphere of the liposome. MTs are shown in green. Kinesins attached to the liposome but not engaged with the MT are shown in magenta. Kinesins engaged with the MT are shown as red and yellow heads with a magenta stalk. A red line represents the attachment between the kinesin tail and the liposome and the length of the line represents the force on that attachment point. Unlike Video S7 where the tug-of-war results in the liposome turning onto the crossing MT, the tug-of-war here is unproductive and leaves the liposome stuck in a position where it cannot proceed, inevitably leading to a termination outcome.

Movie S9. *Simulated liposomes with reduced motor number can terminate in intersections without a tug-of-war.* Video of simulated liposome with 3 liposome-bound kinesins terminating in a 3D MT intersection. The liposome is shown in gold with the blue band indicating the hemisphere of the liposome. MTs are shown in green. Kinesins attached to the liposome but not engaged with the MT are shown in magenta. Kinesins engaged with the MT are shown as red and yellow heads with a magenta stalk. A red line represents the attachment between the kinesin tail and the liposome and the length of the line represents the force on that attachment point. Notice that the liposome terminates in the intersection shortly after reaching and colliding with the crossing MT. The force felt by the motor engaged with the starting MT rises until that motor undergoes force-dependent detachment from the MT, terminating the run.

## References

1. Coy, D.L., W.O. Hancock, M. Wagenbach, and J. Howard. 1999. Kinesin’s tail domain is an inhibitory regulator of the motor domain. Nat Cell Biol. 1:288–292.

2. Vale, R.D. 2003. The Molecular Motor Toolbox for Intracellular Transport. Cell. 112:467– 480.

3. Heaslip, A.T., S.R. Nelson, A.T. Lombardo, S. Beck Previs, J. Armstrong, and D.M. Warshaw. 2014. Cytoskeletal Dependence of Insulin Granule Movement Dynamics in INS-1 Beta-Cells in Response to Glucose. PLoS ONE. 9:e109082.

4. Cai, D., A.D. Hoppe, J.A. Swanson, and K.J. Verhey. 2007. Kinesin-1 structural organization and conformational changes revealed by FRET stoichiometry in live cells. The Journal of Cell Biology. 176:51–63.

5. Hirokawa, N., Y. Noda, Y. Tanaka, and S. Niwa. 2009. Kinesin superfamily motor proteins and intracellular transport. Nat Rev Mol Cell Biol. 10:682–696.

6. Hirokawa, N., and Y. Tanaka. 2015. Kinesin superfamily proteins (KIFs): Various functions and their relevance for important phenomena in life and diseases. Experimental Cell Research. 334:16–25.

7. Fourriere, L., A.J. Jimenez, F. Perez, and G. Boncompain. 2020. The role of microtubules in secretory protein transport. Journal of Cell Science. 133:jcs237016.

8. Barlan, K., and V.I. Gelfand. 2017. Microtubule-Based Transport and the Distribution, Tethering, and Organization of Organelles. Cold Spring Harb Perspect Biol. 9:a025817.

9. Friedman, D.S., and R.D. Vale. 1999. Single-molecule analysis of kinesin motility reveals regulation by the cargo-binding tail domain. Nat Cell Biol. 1:293–297.

10. Reilein, A.R., S.L. Rogers, M.C. Tuma, and V.I. Gelfand. 2001. Regulation of molecular motor proteins. In: International Review of Cytology. Elsevier. pp. 179–238.

11. Verhey, K.J., and J.W. Hammond. 2009. Traffic control: regulation of kinesin motors. Nat Rev Mol Cell Biol. 10:765–777.

12. Ross, J.L., M.Y. Ali, and D.M. Warshaw. 2008. Cargo transport: molecular motors navigate a complex cytoskeleton. Current Opinion in Cell Biology. 20:41–47.

13. Vale, R., T. Reese, and M. Sheetz. 1985. Identification of a novel force-generating protein, kinesin, involved in microtubule-based motility. Cell. 42:39–50.

14. Verhey, K.J., N. Kaul, and V. Soppina. 2011. Kinesin Assembly and Movement in Cells. Annu. Rev. Biophys. 40:267–288.

15. Weijman, J.F., S.K.N. Yadav, K.J. Surridge, J.A. Cross, U. Borucu, J. Mantell, D.N. Woolfson, C. Schaffitzel, and M.P. Dodding. 2022. Molecular architecture of the autoinhibited kinesin-1 lambda particle. Sci. Adv. 8:eabp9660.

16. Chiba, K., K.M. Ori-McKenney, S. Niwa, and R.J. McKenney. 2022. Synergistic autoinhibition and activation mechanisms control kinesin-1 motor activity. Cell Reports. 39:110900.

17. Tan, Z., Y. Yue, F. Leprevost, S. Haynes, V. Basrur, A.I. Nesvizhskii, K.J. Verhey, and M.A. Cianfrocco. 2023. Autoinhibited kinesin-1 adopts a hierarchical folding pattern. eLife. 12:RP86776.

18. Carrington, G., U. Fatima, I. Caramujo, T. Lewis, D. Casas-Mao, and M. Peckham. 2024. A multiscale approach reveals the molecular architecture of the autoinhibited kinesin KIF5A. J Biol Chem. 300:105713.

19. Fu, M., and E.L.F. Holzbaur. 2014. Integrated regulation of motor-driven organelle transport by scaffolding proteins. Trends in Cell Biology. 24:564–574.

20. Bensel, B.M., S.B. Previs, C. Bookwalter, K.M. Trybus, S. Walcott, and D.M. Warshaw. 2024. Kinesin-1-transported liposomes prefer to go straight in 3D microtubule intersections by a mechanism shared by other molecular motors. Proc. Natl. Acad. Sci. U.S.A. 121:e2407330121.

21. Bálint, Š., I. Verdeny Vilanova, Á. Sandoval Álvarez, and M. Lakadamyali. 2013. Correlative live-cell and superresolution microscopy reveals cargo transport dynamics at microtubule intersections. Proc. Natl. Acad. Sci. U.S.A. 110:3375–3380.

22. Lakadamyali, M. 2014. Navigating the cell: how motors overcome roadblocks and traffic jams to efficiently transport cargo. Phys. Chem. Chem. Phys. 16:5907.

23. Osunbayo, O., J. Butterfield, J. Bergman, L. Mershon, V. Rodionov, and M. Vershinin. 2015. Cargo Transport at Microtubule Crossings: Evidence for Prolonged Tug-of-War between Kinesin Motors. Biophysical Journal. 108:1480–1483.

24. Verdeny-Vilanova, I., F. Wehnekamp, N. Mohan, Á.S. Álvarez, J.S. Borbely, J.J. Otterstrom, D.C. Lamb, and M. Lakadamyali. 2017. 3D motion of vesicles along microtubules helps them to circumvent obstacles in cells. Journal of Cell Science. jcs.201178.

25. Lu, H., M.Y. Ali, C.S. Bookwalter, D.M. Warshaw, and K.M. Trybus. 2009. Diffusive Movement of Processive Kinesin-1 on Microtubules. Traffic. 10:1429–1438.

26. Randall, T.S., Y.Y. Yip, D.J. Wallock-Richards, K. Pfisterer, A. Sanger, W. Ficek, R.A. Steiner, A.J. Beavil, M. Parsons, and M.P. Dodding. 2017. A small-molecule activator of kinesin-1 drives remodeling of the microtubule network. Proc Natl Acad Sci U S A. 114:13738–13743.

27. Cronan, J.E. 1990. Biotination of proteins in vivo. A post-translational modification to label, purify, and study proteins. J Biol Chem. 265:10327–10333.

28. Ali, M.Y., H. Lu, C.S. Bookwalter, D.M. Warshaw, and K.M. Trybus. 2008. Myosin V and Kinesin act as tethers to enhance each others’ processivity. Proc. Natl. Acad. Sci. U.S.A. 105:4691–4696.

29. Nelson, S.R., K.M. Trybus, and D.M. Warshaw. 2014. Motor coupling through lipid membranes enhances transport velocities for ensembles of myosin Va. Proc. Natl. Acad. Sci. U.S.A. 111.

30. Lombardo, A.T., S.R. Nelson, M.Y. Ali, G.G. Kennedy, K.M. Trybus, S. Walcott, and D.M. Warshaw. 2017. Myosin Va molecular motors manoeuvre liposome cargo through suspended actin filament intersections in vitro. Nat Commun. 8:15692.

31. Schindelin, J., I. Arganda-Carreras, E. Frise, V. Kaynig, M. Longair, T. Pietzsch, S. Preibisch, C. Rueden, S. Saalfeld, B. Schmid, J.-Y. Tinevez, D.J. White, V. Hartenstein, K. Eliceiri, P. Tomancak, and A. Cardona. 2012. Fiji: an open-source platform for biological-image analysis. Nat Methods. 9:676–682.

32. Lombardo, A.T., S.R. Nelson, G.G. Kennedy, K.M. Trybus, S. Walcott, and D.M. Warshaw. 2019. Myosin Va transport of liposomes in three-dimensional actin networks is modulated by actin filament density, position, and polarity. Proc. Natl. Acad. Sci. U.S.A. 116:8326– 8335.

33. Nayak, C.R., and A.D. Rutenberg. 2011. Quantification of Fluorophore Copy Number from Intrinsic Fluctuations during Fluorescence Photobleaching. Biophysical Journal. 101:2284– 2293.

34. McKenney, R.J., W. Huynh, M.E. Tanenbaum, G. Bhabha, and R.D. Vale. 2014. Activation of cytoplasmic dynein motility by dynactin-cargo adapter complexes. Science. 345:337– 341.

35. Soppina, V., S.R. Norris, A.S. Dizaji, M. Kortus, S. Veatch, M. Peckham, and K.J. Verhey. 2014. Dimerization of mammalian kinesin-3 motors results in superprocessive motion. Proc. Natl. Acad. Sci. U.S.A. 111:5562–5567.

36. Mayr, M.I., M. Storch, J. Howard, and T.U. Mayer. 2011. A Non-Motor Microtubule Binding Site Is Essential for the High Processivity and Mitotic Function of Kinesin-8 Kif18A. PLoS ONE. 6:e27471.

37. Bayerl, T.M., and M. Bloom. 1990. Physical properties of single phospholipid bilayers adsorbed to micro glass beads. A new vesicular model system studied by 2H-nuclear magnetic resonance. Biophysical Journal. 58:357–362.

38. Mornet, S., O. Lambert, E. Duguet, and A. Brisson. 2005. The Formation of Supported Lipid Bilayers on Silica Nanoparticles Revealed by Cryoelectron Microscopy. Nano Lett. 5:281–285.

39. Venables, W.N., and B.D. Ripley. 2002. Modern Applied Statistics with S. Fourth. New York: Springer.

40. Huang, B., W. Wang, M. Bates, and X. Zhuang. 2008. Three-Dimensional Super-Resolution Imaging by Stochastic Optical Reconstruction Microscopy. Science. 319:810–813.

41. Henriques, R., M. Lelek, E.F. Fornasiero, F. Valtorta, C. Zimmer, and M.M. Mhlanga. 2010. QuickPALM: 3D real-time photoactivation nanoscopy image processing in ImageJ. Nat Methods. 7:339–340.

42. Katrukha, E., Jalmar Teeuw, Bmccloin, and J.D. Braber. 2022. ekatrukha/DoM_Utrecht: Detection of Molecules 1.2.5..

43. Bergman, J.P., M.J. Bovyn, F.F. Doval, A. Sharma, M.V. Gudheti, S.P. Gross, J.F. Allard, and M.D. Vershinin. 2018. Cargo navigation across 3D microtubule intersections. Proc. Natl. Acad. Sci. U.S.A. 115:537–542.

44. Bovyn, M., B.R. Janakaloti Narayanareddy, S. Gross, and J. Allard. 2021. Diffusion of kinesin motors on cargo can enhance binding and run lengths during intracellular transport. MBoC. 32:984–994.

45. Sarpangala, N., and A. Gopinathan. 2022. Cargo surface fluidity can reduce inter-motor mechanical interference, promote load-sharing and enhance processivity in teams of molecular motors. PLoS Comput Biol. 18:e1010217.

46. Andreasson, J.O.L., S. Shastry, W.O. Hancock, and S.M. Block. 2015. The Mechanochemical Cycle of Mammalian Kinesin-2 KIF3A/B under Load. Current Biology. 25:1166–1175.

47. Gross, S.P., M. Vershinin, and G.T. Shubeita. 2007. Cargo Transport: Two Motors Are Sometimes Better Than One. Current Biology. 17:R478–R486.

48. Vershinin, M., B.C. Carter, D.S. Razafsky, S.J. King, and S.P. Gross. 2007. Multiple-motor based transport and its regulation by Tau. Proc. Natl. Acad. Sci. U.S.A. 104:87–92.

49. Furuta, K., A. Furuta, Y.Y. Toyoshima, M. Amino, K. Oiwa, and H. Kojima. 2013. Measuring collective transport by defined numbers of processive and nonprocessive kinesin motors. Proc. Natl. Acad. Sci. U.S.A. 110:501–506.

50. Feng, Q., K.J. Mickolajczyk, G.-Y. Chen, and W.O. Hancock. 2018. Motor Reattachment Kinetics Play a Dominant Role in Multimotor-Driven Cargo Transport. Biophysical Journal. 114:400–409.

51. Rogers, A.R., J.W. Driver, P.E. Constantinou, D. Kenneth Jamison, and M.R. Diehl. 2009. Negative interference dominates collective transport of kinesin motors in the absence of load. Phys. Chem. Chem. Phys. 11:4882.

52. Jamison, D.K., J.W. Driver, A.R. Rogers, P.E. Constantinou, and M.R. Diehl. 2010. Two Kinesins Transport Cargo Primarily via the Action of One Motor: Implications for Intracellular Transport. Biophysical Journal. 99:2967–2977.

53. Hendricks, A.G., E.L.F. Holzbaur, and Y.E. Goldman. 2012. Force measurements on cargoes in living cells reveal collective dynamics of microtubule motors. Proc. Natl. Acad. Sci. U.S.A. 109:18447–18452.

54. Visscher, K., M.J. Schnitzer, and S.M. Block. 1999. Single kinesin molecules studied with a molecular force clamp. Nature. 400:184–189.

55. Schnitzer, M.J., K. Visscher, and S.M. Block. 2000. Force production by single kinesin motors. Nat Cell Biol. 2:718–723.

56. Schroeder, H.W., A.G. Hendricks, K. Ikeda, H. Shuman, V. Rodionov, M. Ikebe, Y.E. Goldman, and E.L.F. Holzbaur. 2012. Force-Dependent Detachment of Kinesin-2 Biases Track Switching at Cytoskeletal Filament Intersections. Biophysical Journal. 103:48–58.

57. Angerani, S., E. Lindberg, N. Klena, C.K.E. Bleck, C. Aumeier, and N. Winssinger. 2021. Kinesin-1 activity recorded in living cells with a precipitating dye. Nat Commun. 12:1463.

58. D’Souza, A.I., R. Grover, G.A. Monzon, L. Santen, and S. Diez. 2023. Vesicles driven by dynein and kinesin exhibit directional reversals without regulators. Nat Commun. 14:7532.

59. Gicking, A.M., T.-C. Ma, Q. Feng, R. Jiang, S. Badieyan, M.A. Cianfrocco, and W.O. Hancock. 2022. Kinesin-1, −2, and −3 motors use family-specific mechanochemical strategies to effectively compete with dynein during bidirectional transport. eLife. 11:e82228.

60. Monroy, B.Y., T.C. Tan, J.M. Oclaman, J.S. Han, S. Simó, S. Niwa, D.W. Nowakowski, R.J. McKenney, and K.M. Ori-McKenney. 2020. A Combinatorial MAP Code Dictates Polarized Microtubule Transport. Developmental Cell. 53:60–72.e4.

61. Ferro, L.S., Q. Fang, L. Eshun-Wilson, J. Fernandes, A. Jack, D.P. Farrell, M. Golcuk, T. Huijben, K. Costa, M. Gur, F. DiMaio, E. Nogales, and A. Yildiz. 2022. Structural and functional insight into regulation of kinesin-1 by microtubule-associated protein MAP7. Science. 375:326–331.

62. Soppina, V., and K.J. Verhey. 2014. The family-specific K-loop influences the microtubule on-rate but not the superprocessivity of kinesin-3 motors. MBoC. 25:2161–2170.

63. Gilbert, S.P., S. Guzik-Lendrum, and I. Rayment. 2018. Kinesin-2 motors: Kinetics and biophysics. Journal of Biological Chemistry. 293:4510–4518.

