## Supplemental Materials for "Kinesin-1 Autoinhibition Tunes Cargo Transport by Motor Ensembles"

Supplemental Figures

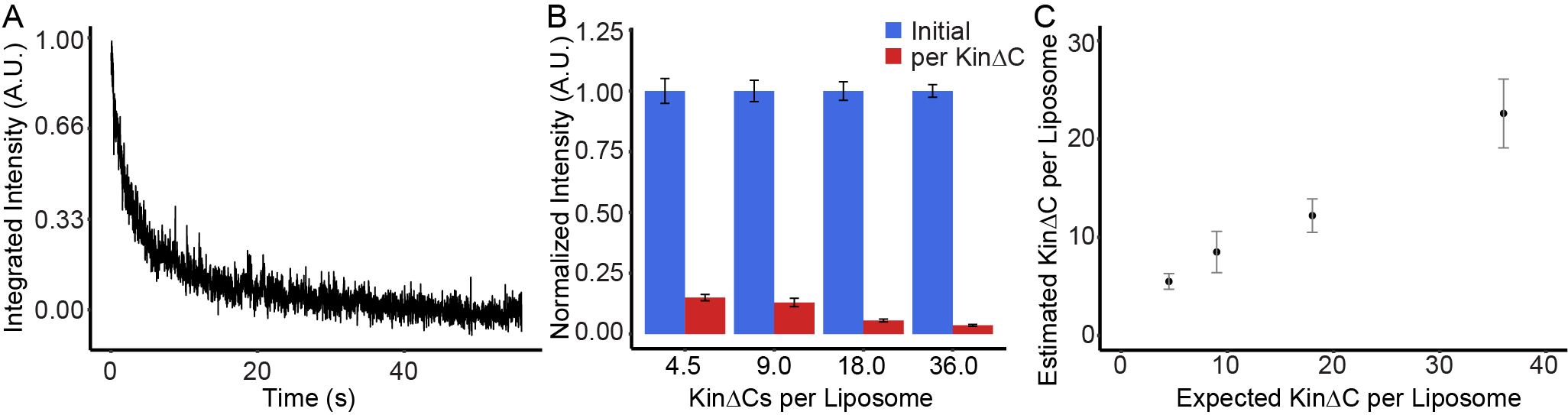

Fig. S1. *Motor Density Estimation by Photobleaching of KinΔC-YFP.* (A) Example fluorescence decay curve recorded from a liposome incubated with a 9-fold excess of KinΔC-YFP. (B) Bar plot of normalized initial intensity (blue) and normalized molecular intensity of KinΔC-YFP (red) as determined for each incubation ratio used to build a standard curve. (C) Plot of estimated KinΔC per liposome versus expected KinΔC per liposome, i.e., the incubation ratio. See Methods for a detailed explanation of the technique.

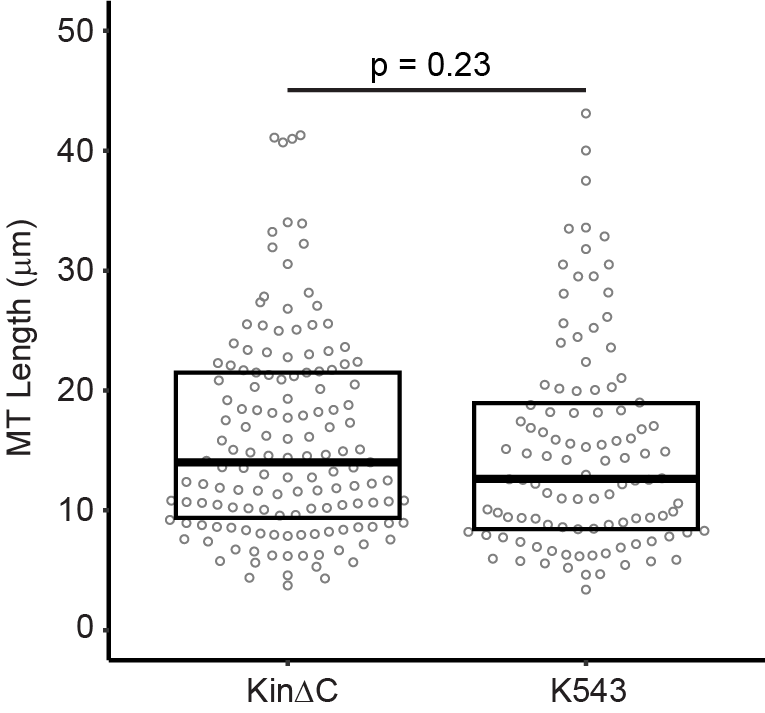

Fig. S2. *MT length distributions for KinΔC and K543 multi-motor motility assays.* Dot plot with overlaid box plot of the lengths of the MTs used in run length analysis of KinΔC and K543 single molecules and ensembles. P-Value is reported from a Kruskal-Wallace rank sum test. N = 144 MTs (KinΔC) and 106 MTs (K543) from 3 independent MT preparations each. K543 data are replotted from Bensel et al., 2024.

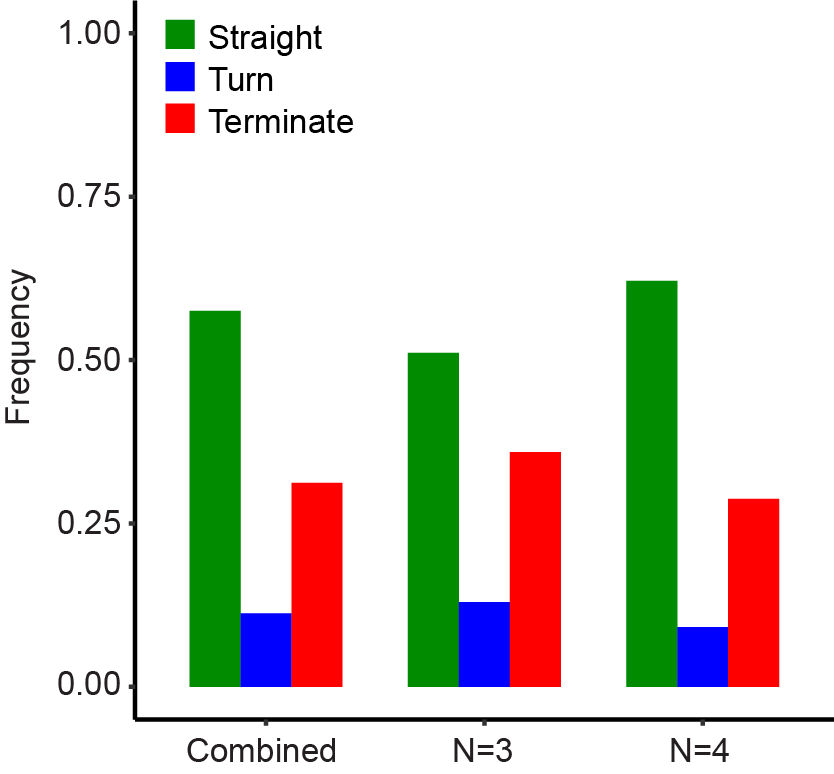

Fig. S3. *Simulated 3D intersection outcomes for a mixture of 3 and 4 bound kinesins, 3 kinesins alone, and 4 kinesins alone.* Bar graph of outcome frequency for mechanistic models with reduced motor number. Because a 3-fold reduction from 10 is 3.33, we performed simulations with either 3 or 4 motors bound to the liposome and then combined the results. Here we show the results split out versus to combination. For N=3 case, the data come from 54 simulated events, while the data for the N=4 case comes from 57 simulated events.

| **Comparison** | **Run Length** | **End Events** | **Velocity** | **F_Detach_** | **Landing Rate** |
| --- | --- | --- | --- | --- | --- |
| *1 KinΔC vs 1 K543* | 1.00 | 0.62 | 0.71 | 0.15 | 0.01 |
| *5 KinΔC vs 5 K543* | 0.042 | 0.12 | 0.0005 | NA | NA |
| *10 KinΔC vs 10 K543* | 0.046 | 0.0003 | 1 | NA | NA |
| *20 KinΔC vs 20 K543* | 0.000019 | 0.0001 | 0.00003 | 1.32 x 10^-9^ | NA |
| *KinΔC* *± Kinesore* | NA | NA | NA | 3.11 x 10^-6^ | 0.0003 |
| *K543* *± Kinesore* | NA | NA | NA | NA | 0.89 |
| *KinΔC+Kinesore vs K543* | NA | NA | NA | NA | 0.90 |
| *1 KinΔC vs 5 KinΔC* | NA | NA | 1.75 x 10^-10^ | NA | NA |
| *1 KinΔC vs 10 KinΔC* | NA | NA | 2.02 x 10^-21^ | NA | NA |
| *1 KinΔC vs 20 KinΔC* | NA | NA | 1.00 x 10^-34^ | NA | NA |
| *Active Model vs Fast k_+_/k_-_* | 0.0156 | NA | 0.383 | NA | NA |
| *Active Model vs Slow k_+_/k_-_* | 0.0411 | NA | 0.499 | NA | NA |

**Table S1:** *Statistical Analysis of Differences Between KinΔC and K543 Control.* Related to Figures 2 and 3. Run length, velocity, and landing rate comparisons were made using a Kuskal-Wallis with Dunn’s post-hoc test using the Benjamin-Yuketieli adjustment. End Event Frequencies were compared using a two proportions Z test. Detachment force distributions were compared using an exact two-sample Kolmogorov-Smirnov test. p-Values less than 0.05 are considered statistically significant.
